# Problems using ordinal traits with continuous measures of functional diversity

**DOI:** 10.1101/2021.11.02.466687

**Authors:** Matt Davis

## Abstract

Continuous indices of functional diversity are popular in studies examining community structure and ecosystem function across a wide range of subfields from paleontology to range management. These indices were designed to replace the use of more arbitrary, discrete functional groups or guilds; however, the effect of typical methodological decisions on these new continuous measures has not been fully investigated. To test the effect of using ordinal traits in functional diversity analysis, I first calculated relative functional diversity index values in real plant communities with real continuous trait data and Euclidean distances. I then compared these original values to “treatment” functional diversity index values obtained by discretizing the trait data and using Gower’s distance. Agreement between original and treatment values was highly unpredictable and often abysmal. Small methodological choices, such as whether to treat a functional trait as continuous (mm) or ordinal (“small”, “medium”, “large”), could completely change a perceived functional diversity relationship along an environmental gradient. Some parameter combinations returned results that were no better than random noise. Because simple methodological choices can have such a large impact on continuous functional diversity indices, it is ambiguous whether analyses using ordinal traits are actually measuring an underlying functional diversity relationship between communities or just reflecting the arbitrary parameter choices of researchers.

## Introduction

Although often described as a burgeoning new subfield (Petchey and Gaston 2006), or even a bandwagon (McGill 2015), the study of functional trait diversity, the range and value of species or organismal traits that effect ecosystem functioning (Tilman 2001), has long been an integral part of ecological theory and practice previously investigated through morphospace or functional groups and guilds (e.g. Holmes et al. 1979, Van Valkenburgh 1988). Despite mixed evidence (Stevens and Tello 2014), functional diversity is thought to represent a unique aspect of biodiversity (Devictor et al. 2010) that could potentially explain ecosystem function, species coexistence, and community assembly better than taxonomic richness or phylogenetic diversity alone (Tilman et al. 1997, Hulot et al. 2000, Díaz and Cabido 2001, Loreau et al. 2001, Heemsbergen et al. 2004, Hooper et al. 2005, Mason et al. 2005, Pavoine and Bonnan 2011). Similar to phylogenetic diversity, which moved from subjectively defined taxonomic units to increasingly sophisticated statistical methods of tree discovery (Felsenstein 2004, Redding et al. 2014), functional diversity has changed from the census of discrete, arbitrarily defined functional guilds (e.g. large carnivore, medium carnivore) (Fonseca and Ganade 2001) to complex measures of multidimensional functional space (Mouchet et al. 2010, Schleuter et al. 2010) that can be used to model future structural changes in ecosystems (Barbet-Massin and Jetz 2015) or triage species endangered by climate change (Buisson et al. 2013).

Central to this transformation was the view that a priori discrete functional groups, although frequently used, were not based on objective criteria (Petchey et al. 2009), overestimated redundancy by lumping together heterogeneous species (Fonseca and Ganade 2001), and often provided no predictive power over random classifications (Wright et al. 2005). New measures of functional diversity had to be continuous and relatively independent of arbitrary researcher decisions (Petchey et al. 2004). Though others had used both multidimensional functional space and functional dendrograms before to examine community structure (e.g. Holmes et al. 1979), the theoretical framework for these modern indices was laid by Rosenfeld (2002) who formally defined functional space as a hypervolume delineated by axes of functional traits or processes (e.g. CO2 production, N fixation, etc.) analogous to Hutchinson’s (1957) n-dimensional niche defined by environmental axes. Using this functional space, Villéger et al. (2008) developed a general methodology for calculating continuous functional diversity indices that was expanded on and operationalized by Laliberté and Legendre (2010) into the now popular R software package “FD”. First, researchers measure ostensibly functionally important traits of taxa (typically species) and after standardization, use these traits to generate a species-species dissimilarity matrix. These interspecies distances are subjected to a principle coordinates analysis (PCoA) that places species in a multidimensional Euclidean trait space where a range of functional index values can be calculated on the species’ new traits (the PCoA axes). This methodological framework is also used to construct the popular dendrogram-based functional richness index (FD) developed by Petchey and Gaston (2002, 2007) and modified by Podani and Scherma (2006) except that, instead of PCoA, the species-species distance matrix is run through a sorting algorithm such as unweighted pair-group clustering method using arithmetic averages (UPGMA) to generate a functional tree with species as tips.

Calculating these Euclidean distances between species is straightforward with continuous traits measured on interval and ratio-scales such as body temperature (ºC) or leaf area (mm2), respectively, but some continuous traits such as flowering time (circular variable), or percentage of diet from different feeding categories (fuzzy variable) cannot immediately be compared using Euclidean distances without some kind of transformation. Many traits are discrete, such as presence or absence of spines (binary variable), growth form (multistate nominal variable), or height class (ordinal variable), and aside from binary variables, they cannot be readily converted into Euclidean distances. This is why all developers of this common functional diversity methodological framework (Podani and Schmera 2006, Petchey and Gaston 2007, Villéger et al. 2008, Laliberté and Legendre 2010) suggest that researchers use Gower’s general coefficient of similarity (Gower 1971) to calculate distances between taxa when considering variables of mixed types.

Gower originally developed his coefficient for quantitative taxonomy (Pavoine et al. 2009) but its Euclidean properties and adaptability made it widely popular and useful in fields as disparate as retail science (Churchill et al. 1975) and dermatology (Katrina et al. 1985). Gower calculated the similarity *S*_*jk*_ of two taxa (typically species) *j* and *k* as,

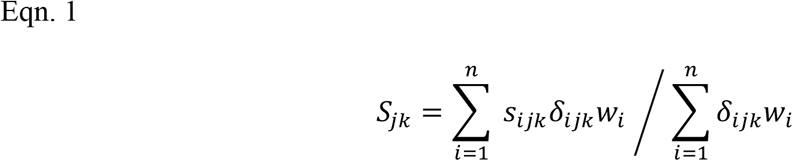

where *n* is the number of traits examined, *w*_*i*_ is the variable weight, *s*_*ijk*_ is the similarity (ranging from 0-1) between taxa *j* and *k* calculated on trait *i*, and *δ*_*i*_ = 0 if the *i*th trait value is missing for one or both taxa and *δ*_*i*_ = 1 if both taxa posses the value and can be compared. Gower defined the similarity *s*_*ijk*_ individually for continuous, nominal, and binary variables (Gower 1971), with later authors defining extensions for ordinal variables (Podani 1999); multichoice nominal and associated-binary variables (Podani and Schmera 2007); and circular, fuzzy, and correlated-proportion variables (Pavoine et al. 2009). In the simplest case where there are no missing values and all traits are weighted equally, Gower’s coefficient is the average of these different similarities:

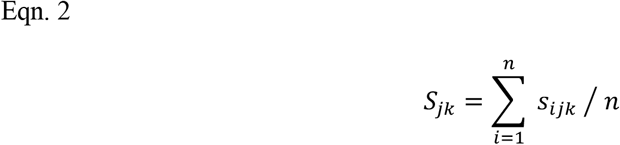

For the rest of this paper, Gower’s coefficient will be treated as a distance metric *D*_*jk*_ = 1 − *S*_*jk*_.

Functional diversity studies implicitly assume that there is some true quantity “functional diversity” that measures the distribution of organisms’ contributions to ecosystem functioning and that this quantity can be estimated by indices calculated from several measurable traits of taxa. However, if methodological choices such as the inclusion of categorical traits requiring Gower’s distance greatly affect relative functional index values between communities, it is doubtful that these indices actually estimate true functional diversity. Continuous measures of functional diversity could suffer from the same types of arbitrary decisions by researchers as the functional guilds they were meant to replace.

Although there is no universally “correct” distance metric, we often intuitively conceptualize functional niche space as a realm measurable by Euclidean distances (**Figure 1**). Hutchinson (Hutchinson 1957) originally defined his n-dimensional hypervolume by environmental variables, “…which can be measured along ordinary rectangular coordinates” but admitted that this unrealistic assumption was a limitation of his concept. Petchey and Gaston (2007) suggest that this distinction may be trivial in practice as they arrived at very similar functional index values whether using Gower’s distance or (incorrectly) using Euclidean distance on datasets with mixed variable types. The users’ manual for the FDiversity software package (Casanoves et al. 2008) also advises that using Gower’s distance may not change results. However, it is clear that using different variable types, and by extension different distance metrics, could produce large downstream changes among analyses. Different variable types lead to specific shapes of point clouds in multidimensional space. Continuous variables produce a continuous line of points but circular variables lead to arcs or circles in space and nominal variables separate points into distinct groups located at the vertices of regular polygons (Pavoine et al. 2009).

**Figure 1.**
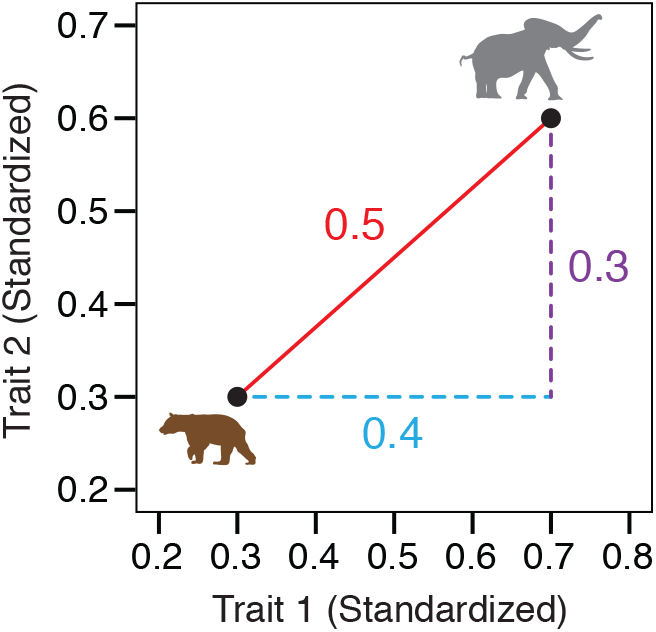
Given a niche space defined by *n* functional traits (standardized by their respective ranges), the difference between two taxa, say a bear and an elephant, is often conceptualized as the Euclidean straight-line distance between them (i.e., 0.5). Gower’s distance however, would take the average of the difference in each functional trait (i.e. (0.4 + 0.3) / 2 = 0.35).

Both Gower’s and Euclidean distances can handle continuous and binary variables, but to measure the impact of using other types of discrete traits, and by extension Gower’s distance, we must consider ordinal characters. Nominal traits have no equivalent in Euclidean space: there are no lines on a ruler for “wind dispersed” and “animal dispersed” by which to measure different seeds. However, ordinal traits such as “small”, “medium”, and “large” typically represent some underlying, measurable continuous distribution for which we lack complete data. By taking a known continuous distribution and discretizing it into ordered levels, we can investigate how inclusion of discrete characters alters relative functional index values between communities. If these indices are accurately estimating some true underlying value of functional diversity, continuous traits and ordinal traits reflecting the same distribution should produce similar results. If index values are wildly different, then we must admit that some measures of functional diversity reflect nothing more than arbitrary methodological choices.

## Methods

In order to test the effect of discretization and Gower’s distance on functional diversity values, I first calculated functional diversity indices with continuous trait data and Euclidean distances following the standard methodology of Laliberté and Legendre (2010). For simplicity, these data, and results generated from them are hereafter referred to as the “original” data/distances/values. I then artificially discretized those continuous trait data and reran analyses using several versions of Gower’s distance. For simplicity, those data and results are referred to as the “treatment” data/distances/values. By comparing relative functional diversity index values between the original and treatment methods, we can see the impact that discrete traits and Gower’s distance have on functional diversity analyses (**Figure 2**).

**Figure 2.**
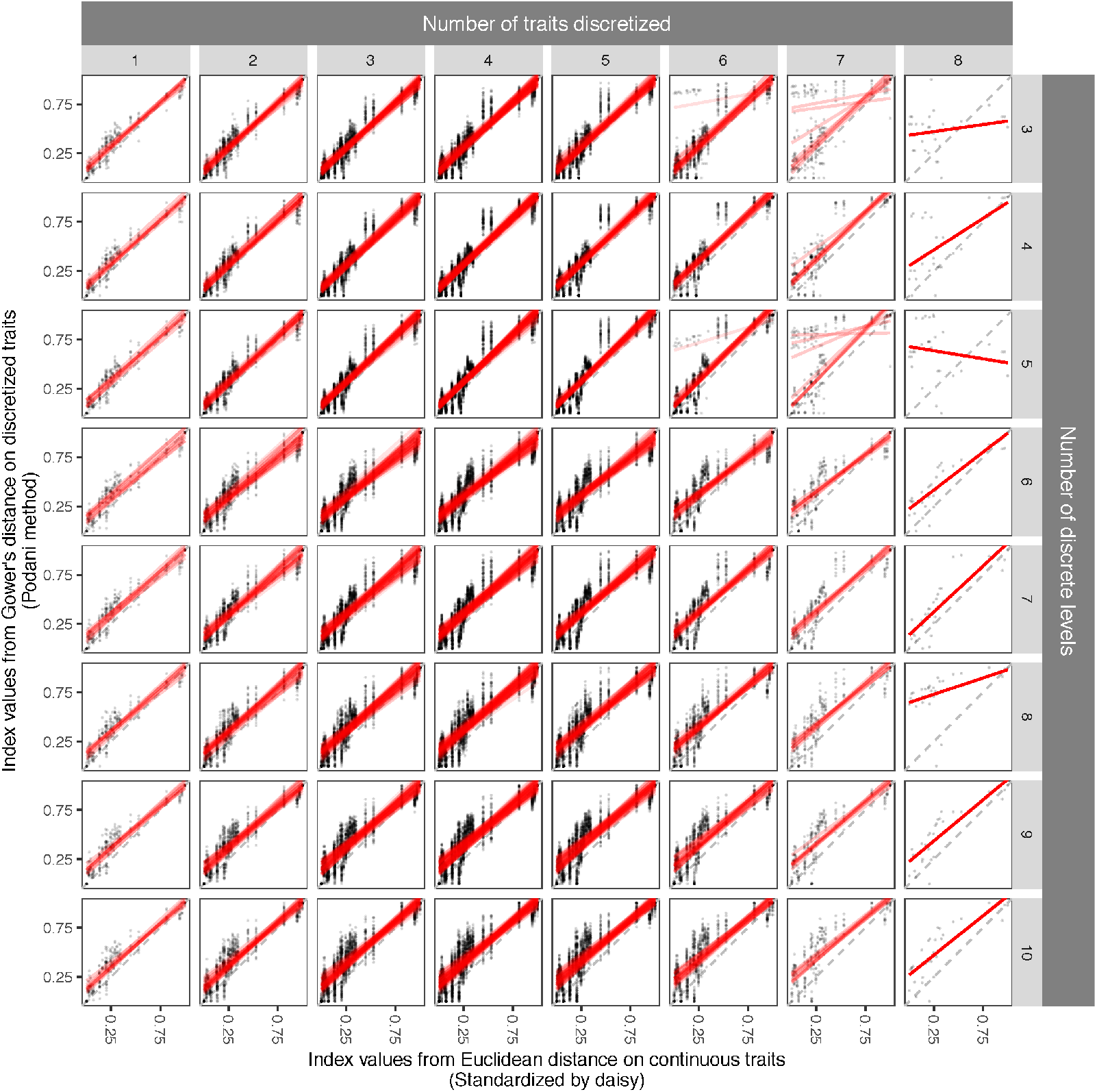
Relative values for functional evenness (FEve) differ between original and treatment versions of the Tussock dataset. Each point represents the relative FEve of a plant community (in this case, experimental plots in New Zealand). On the x-axis, relative FEve values were derived from Euclidean distances between species with original continuous trait data standardized by the daisy R function. On the y-axis, relative FEve values were derived from Gower’s distances between species with trait data artificially discretized into ordered levels. The Podani method was used to calculate Gower’s distance on these treatment data. Dashed grey guidelines mark a 1:1 relationship between the original and treatment data that points would fall on if using discrete characters and different distance metrics had no effect on functional diversity values. Red lines are linear model best fits to the data included only to guide the eye. Most parameter combinations have more than one red line because there are multiple ways to discretize a subset of eight traits (e.g. the two discretized traits could be the 1st and 2nd traits, or the 1st and 3rd traits, or the 2nd and 3rd traits, etc.).

I selected two datasets from the literature for simulating discrete values. The “Tussock” dataset (Laliberté et al. 2008) represents a 2007 census of a series of experimental plots in New Zealand measuring the effect of grazing and fertilizer on plant functional diversity. The “Roadside” dataset (Thompson et al. 2010) represents a series of small quadrats measuring plant species richness every year between 1958 and 2003 along a stretch of road in England. Datasets were trimmed to only continuous traits and any species missing data were removed. Percentage occupancy data were converted to presence/absence by counting all species that occurred in a site. After data conditioning, the Tussock dataset had 41 species with eight traits in 30 communities. The Roadside dataset had 75 species with seven traits in 325 communities (plots measured at different time points). The use of these data should in no way be viewed as a critique on their validity, rather the opposite. I used them because they considered a large number of quantitative traits and their authors graciously made them available.

To create the original functional diversity index values to compare to the discretized treatment data, I first took the Euclidean distance between species based on their continuous traits. Trait values must be standardized so results are not driven by units of measurement (Lepš et al. 2006). I tested the effects of two popular methods of standardization: the defaults for calculating Euclidean and Gower’s distance in R version 3.2.2 (R Development Core Team 2015). In one set of simulations, continuous traits were standardized by the distance function daisy in the R package “cluster” (Maechler et al. 2015) that subtracts the mean of the trait from each value and then divides by the original average distance to the mean for that trait. In the other set of simulations, continuous trait values were divided by trait ranges. Gower (1971) argued that standardizing by trait range made more sense as it could be easily measured on the sample or calculated from theoretical limits (e.g. the minimum and maximum body mass possible for mammals) and it was more immediately interpretable compared to the standard deviation.

Continuous data were discretized into three to ten ordered levels either along absolute splits where the trait range was equally divided or by quantile splits where an equal number of species were included in each level. Although continuous data can be discretized into two ordered levels, in practice they would more likely be dichotomized into binary variables and treated as such. Note that because R collapses levels unoccupied by data points, the given number of levels should be taken as a maximum. For example, a trait split equally along its range into ordered levels 1, 2, 3, and 4 could be collapsed into ordered levels 1, 2, and 3 (formerly 4) if no species occupied the original level 3. Functional trait data were discretized one trait at a time until all traits were discretized into ordered levels. Each possible combination of traits was considered (e.g. the two discretized traits could be the 1st and 2nd traits, or the 1st and 3rd traits, or the 2nd and 3rd traits, etc.). It may seem unrealistic to have a functional dataset where the majority of traits are ordinal or where ordinal traits take more than a few levels but both scenarios are frequent in the literature. For example, 19 out of the 20 functional traits used by Shütz and Schulze (2015) for urban birds were ordinal. In the standardized list of plant traits (Cornelissen et al. 2003), flammability is an ordinal trait with 41 levels.

There are several different ways to calculate dissimilarities with ordinal traits (**Table 1**). Gower (1971) originally lacked a formal measure for such traits, treating ordinal scores as continuous data by taking the Manhattan distance standardized by the range of values (**Table 1**, Continuous), although one could also take the Euclidean distance of these values (**Table 1**, Euclidean). Both methods are mathematically dubious; technically, the differences between ordinal traits are not interpretable (Podani 1999). Sneath and Sokal (1973) proposed that ordinal data should be treated as unranked categories despite a potential loss of information (**Table 1**, Categorical). Podani (1999) created two formal extensions of Gower’s distance. One, metric, took the Manhattan distance between ranks and standardized by the range of ranks (**Table 1**, Metric). The other, nonmetric, calculated the number of interchanges in taxon rank order to move a species from one rank to another and standardized by the total number of interchanges possible (**Table 1**, Podani). This is likely the most commonly used method as it is the default for the popular R program dbFD used in the functional diversity analysis package “FD” (Laliberté and Legendre 2010). Pavoine et al. (2009) proposed another measure analogous to Podani’s (1999) metric equation by taking the Euclidean distance of ranks standardized by the range of ranks, although they took the square root of any resulting distances to lend them Euclidean properties (Gower 1971) (**Table 1**, Pavoine). During this study, I discovered a bug in the code for the Gower’s distance function dist.ktab (Pavoine et al. 2009) in the R package “ade4” (Chessel et al. 2004) that can effectively randomize a matrix of ordinal trait data. To avoid this bug, all ordinal traits should be designated as “numeric” class variables in R, not “ordered” “factor” class as one would assume. Any study using this function in “ade4” version 1.7-4 or earlier, before a clarifying warning was added, should be reevaluated.

**Table 1.**
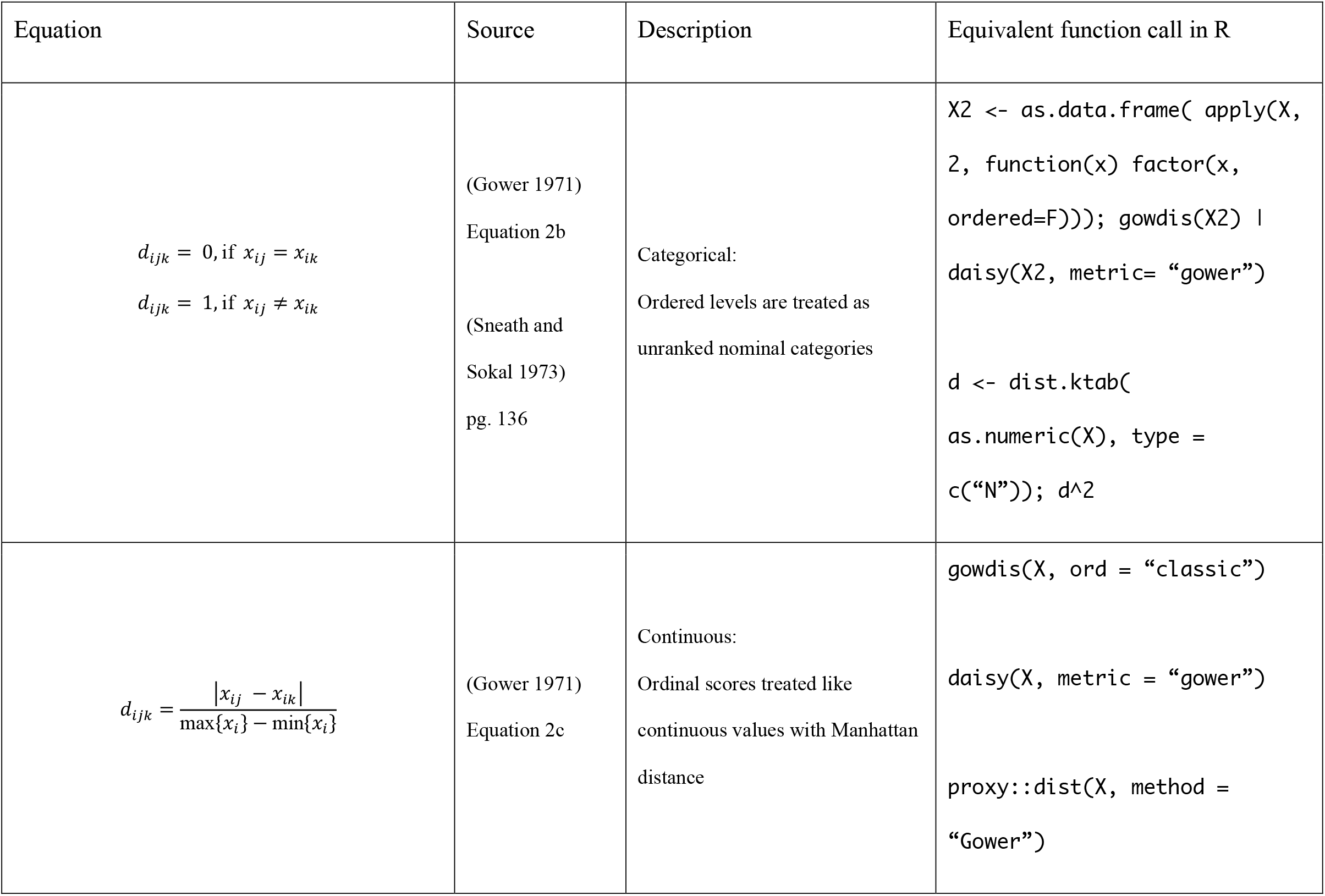
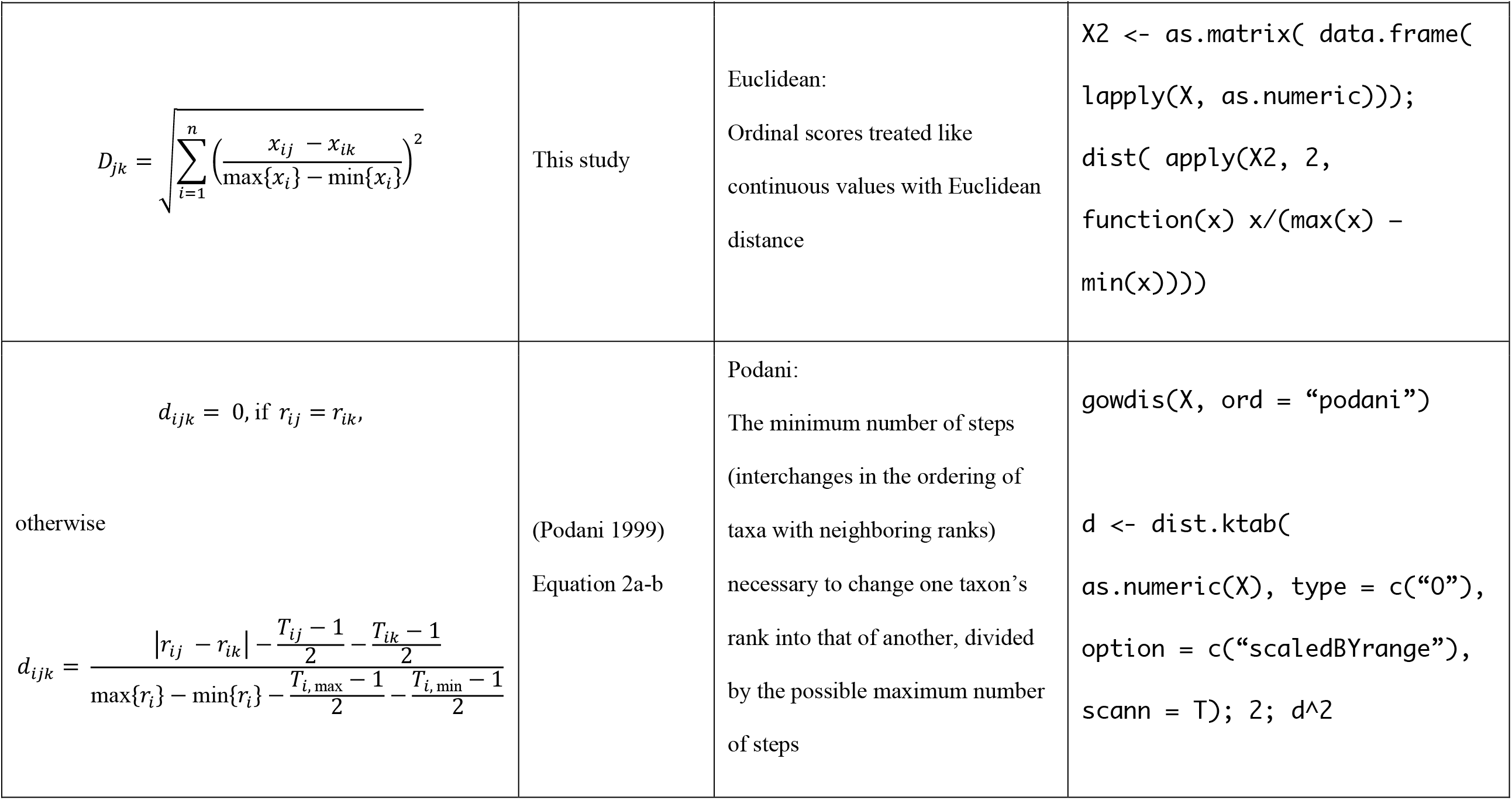
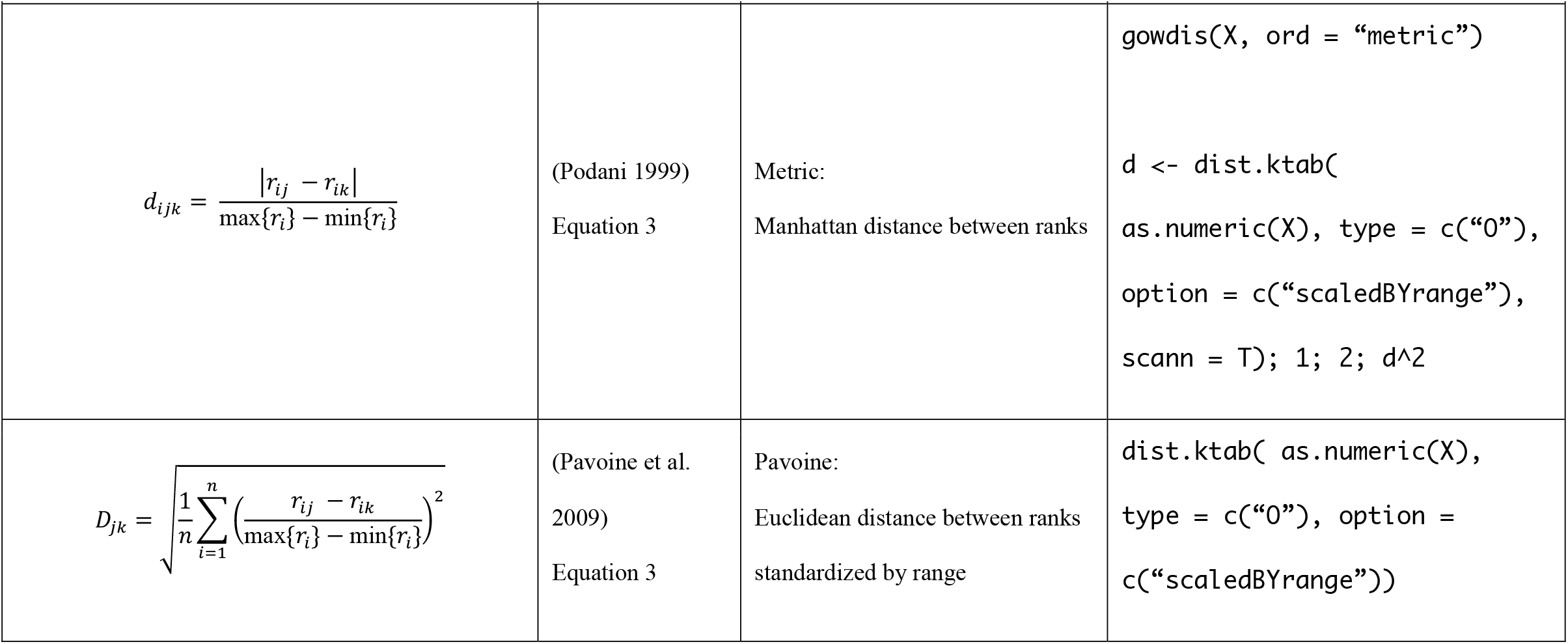
Different methods for calculating differences with ordinal traits. Let X be a matrix containing *m* rows (taxa, typically species or individuals) and *n* columns (traits, characters, variables) of ordinal data. Further, let *i* denote the index of a trait out of the *n* traits considered. Let *x*_*ij*_ be the value taken by this trait for taxon *j*. Let *d*_*ijk*_ be the difference between taxa *j* and *k* as measured by trait *i* with *D*_*jk*_ being the total difference between the two taxa measured on all *n* traits. Let *r*_*ij*_ be the rank of taxon’s *j* ordinal score among all scores of trait *i*. Let *T*_*ij*_ be the number of taxa that have the same rank for trait *i* as taxon *j* (including *j* in this count). Let *T*_*i*, max_ be the number of taxa that have the maximum rank (max{*r*_*i*_}) for trait *i*.

**Table 2.** Popular continuous indices for measuring functional diversity.

| Index | Name | Definition | Low value | High Value | Source |
| --- | --- | --- | --- | --- | --- |
| FD | Functional diversity | Sum of branch lengths on a functional dendrogram needed to connect all taxa in a community | 0: Taxa are identical in functional traits | Unbounded:<br>Taxa are far apart in trait space | (Petchey and Gaston 2007) |
| FRic | Functional richness | Volume of minimum convex hull in trait space enclosing all taxa in a community | 0: Taxa are identical in functional traits | Unbounded:<br>Taxa are far apart in trait space | (Villéger et al. 2008) |
| FEve | Functional evenness | Regularity of taxa along a minimum spanning tree in trait space | 0: Taxa are unevenly distributed throughout trait space | 1: Taxa are evenly distributed throughout trait space | (Villéger et al. 2008) |
| FDiv | Functional divergence | Average deviance from the mean distance to the centroid of trait space | 0: Taxa are close to center of trait space | 1: Taxa are at trait extremes | (Villéger et al. 2008) |
| FDis | Functional dispersal | Mean distance to centroid of trait space | 0: Species are clustered around centroid | Unbounded:<br>Taxa are far from center of trait space | (Laliberté and Legendre 2010) |
| RaoQ | Rao's<br>quadratic<br>entropy | Half the average squared<br>functional distance<br>between two randomly<br>selected individuals with<br>replacement | 0: Taxa are closely<br>packed in trait<br>space | Unbounded:<br>Taxa are far from<br>each other | (Champely<br>and Chessel<br>2002) |

Species-species distance matrices were calculated by Euclidean distances on the original continuous traits or Gower’s distances on the discretized traits using the methods for ordinal data described above. Euclidean distances were also calculated on discretized traits as though they were continuous (**Table 1**, Euclidean). Both original and treatment species-species distance matrices were used to calculate six popular indices of functional diversity (**Figure 2**). With the exception of Petchey and Gaston’s (2007) modified dendrogram-based functional richness (FD), they can all be generated with the R package “FD” (Laliberté and Legendre 2010). FD (Petchey and Gaston 2007) and FRic (Villéger et al. 2008) measure the volume of functional trait space occupied. FDiv (Villéger et al. 2008) and FDis (Laliberté and Legendre 2010) measure species distributions relative to the center of trait space. RaoQ (Botta-Dukát 2005) measures how closely species are packed into trait space, and FEve (Villéger et al. 2008) measures how regularly they are packed.

To test how the inclusion of discrete characters affected relative functional diversity values, I calculated the root mean squared error (RMSE) between original community functional diversity index values and treatment index values (**Figure 3**). Unfortunately, a major shortcoming of functional diversity analyses is that raw index values cannot typically be compared between studies. They are informative only in their relative value within a dataset. For this reason, all indices were standardized by the range of community values before being compared to each other. This generates a conservative estimate of error as this standardization increases the likelihood of a 1:1 relationship between original index values and treatment index values compared to some typical ways of standardization, such as making FRic or FD community values proportional to the total convex hull volume or tree length of all species in the dataset, respectively. However, this is probably the only fair way to compare errors between indices, as given other standardization methods, different indices will tend to occupy different proportions of the 0-1 domain. For example, if average community species richness is low compared to regional richness, standardizing by the regional convex hull volume will always produce low community FRic values, and by extension, low RMSE between actual and modeled values.

**Figure 3.**
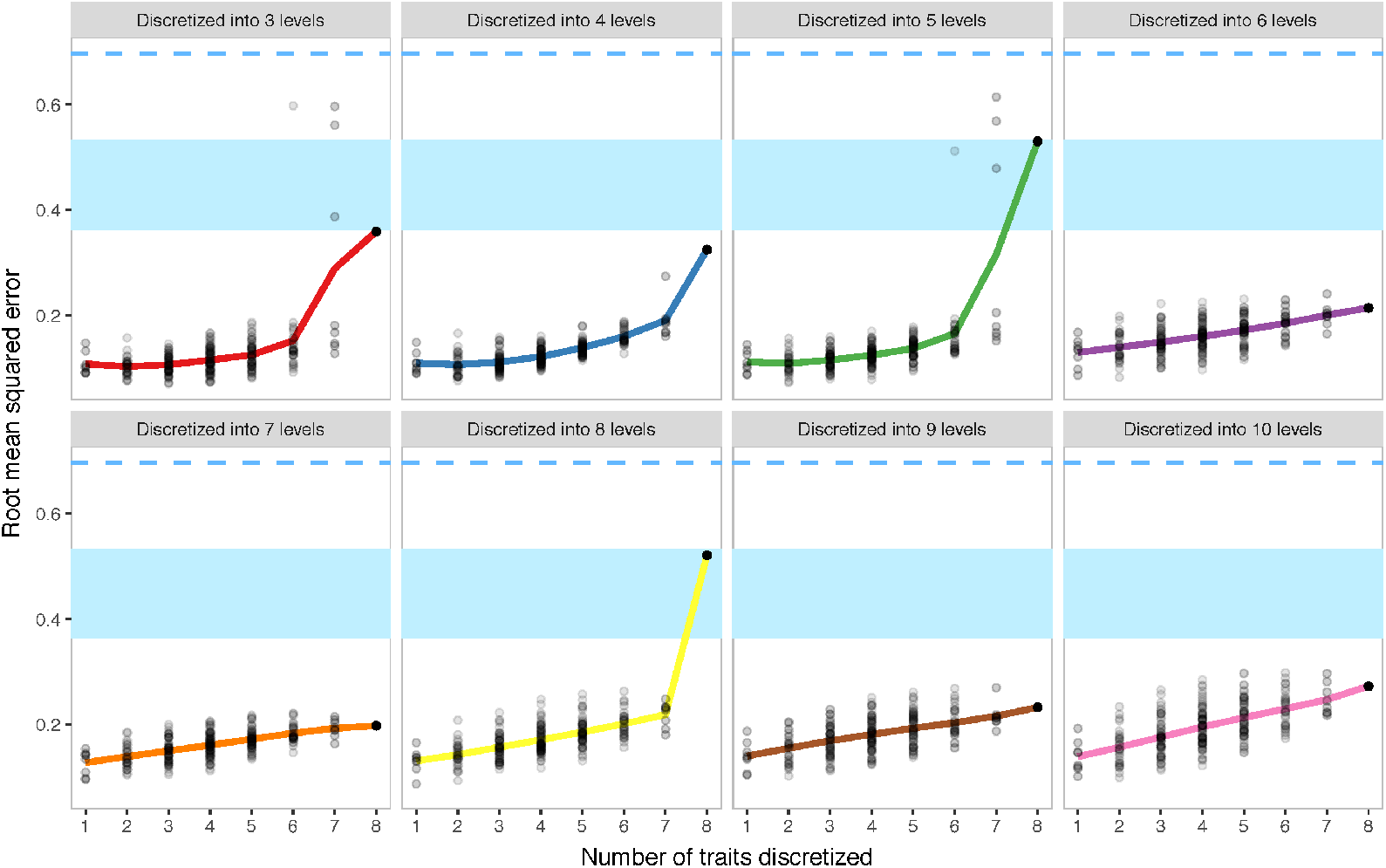
This figure summarizes **Figure 2**; each point represents the root mean squared error (RMSE) between the original functional evenness (FEve) values and one parameter combination of the treatment FEve values. Each colored line connects the means of each parameter combination. The blue dashed line represents the RMSE of a perfect negative correlation to the original FEve values and the blue box represents the 95% confidence interval of the RMSE between the original FEve values and random values from a uniform distribution.

For perspective, these RMSE values were compared to RMSE values obtained from perfect negative correlations to the original indices and the 95% confidence intervals of 100,000 simulations of random values compared to the original indices (**Figure 3, Figure 4**). Note that the confidence intervals for random values are also very conservative. Depending on the underlying structure of the original data, it is possible to find distributions of random values (e.g. drawn from a normal distribution) that are statistically unrelated to the original indices, but still return much lower RMSE values than samples drawn from the uniform distribution bounded between 0 and 1 that was used to construct the confidence intervals. To determine the marginal effect of each parameter on RMSE (**Figure 5, Figure A 1**), I used the partialPlot function on a model generated from 1,001 regression trees in the R package randomForest (Liaw and Wiener 2002) under default parameters leaving a random 30% of the data out for testing. The importance of different parameters for determining RMSE was found by measuring the decrease in the residual sum of squared error in the regression tree when it was split on a given parameter. This value was averaged across all 1,001 trees in the random forest.

**Figure 4.**
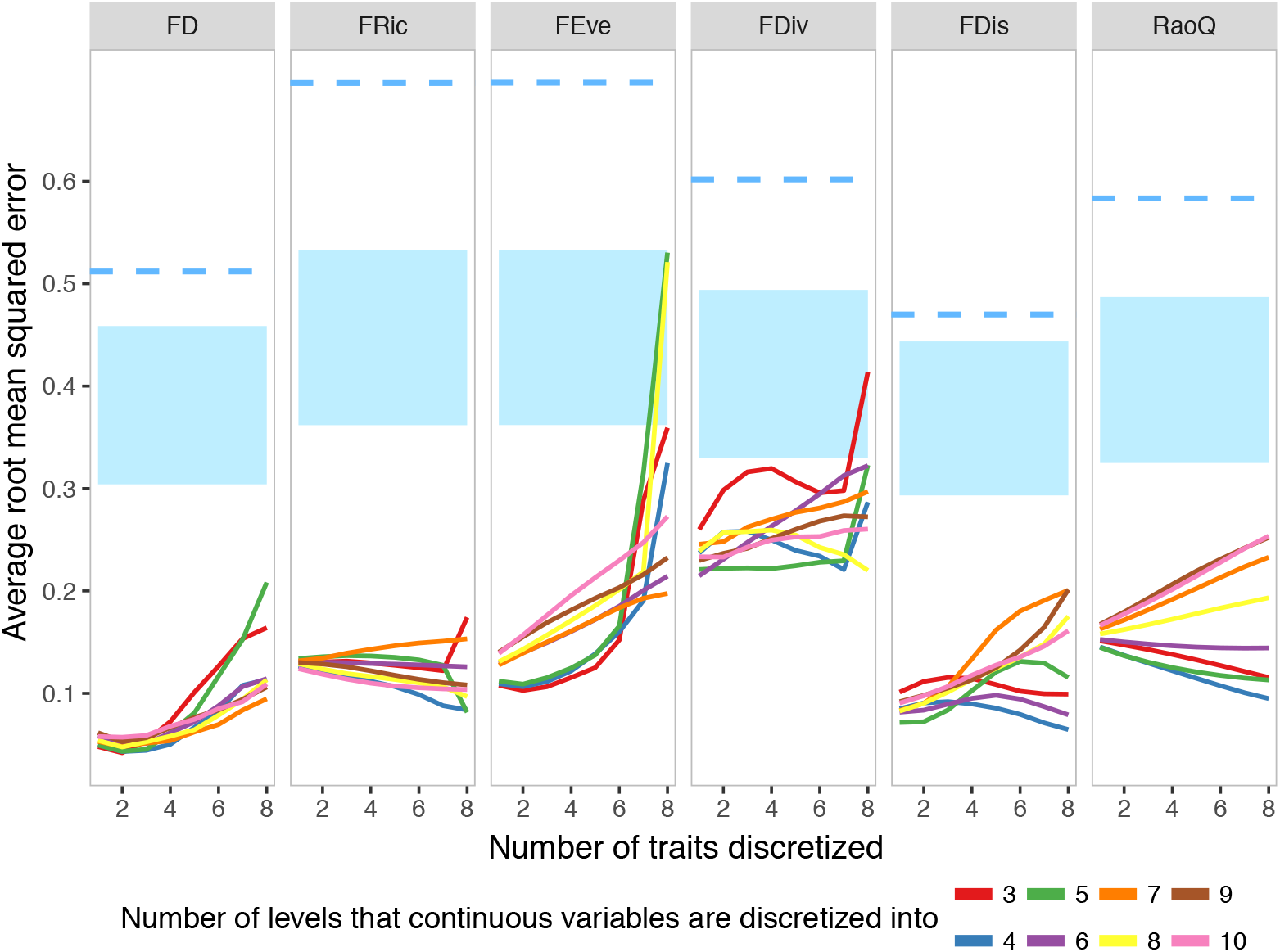
This figure summarizes the root mean squared errors (RMSE) between original functional diversity index values generated from continuous data and the values obtained from different parameter combinations of Gower’s distance using the Podani method on artificially discretized treatment data. Each colored line represents the average RMSE for a particular parameter combination and matches the coloration used in **Figure 3**. The blue dashed lines represent the RMSE of perfect negative correlations to the original functional diversity index values and the blue boxes represent the 95% confidence intervals of the RMSE between the original index values and random values from uniform distributions as in **Figure 3**.

**Figure 5.**
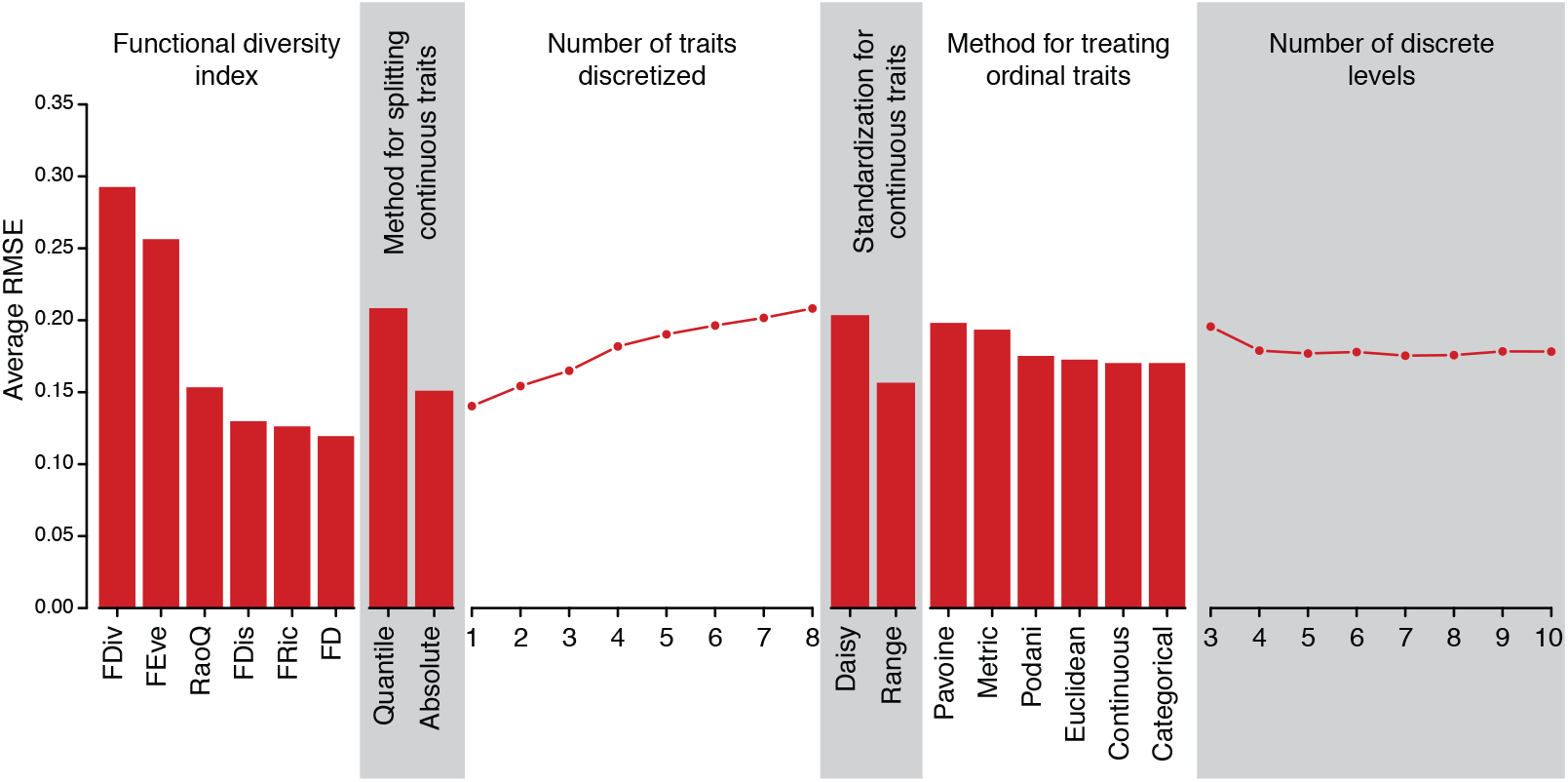
The importance of individual parameters in determining how closely treatment functional diversity values match original index values in the Tussock dataset. The y-axis shows the average root mean squared error (RMSE) between original and treatment values expected from that parameter holding all other parameters constant.

## Results

In general, the inclusion of discrete traits and the use of Gower’s distance had a large effect on functional diversity analyses, although the severity of this effect was highly variable and not always expected. The specific effects of each parameter combination are likely to vary from dataset to dataset, so readers are encouraged to focus on the general qualitative results, rather than the exact values, found here. The model generated from the random forest fit the data well, predicting the 30% left out for testing with a pseudo R^2^ of ∼0.93 for both the Tussock and Roadside datasets. The functional diversity index used was the most important parameter for predicting RMSE, while number of discrete levels was the least important (**Figure 5, Figure A 1**). Within each parameter group, individual parameters varied in how much they affected functional diversity values, although some indices performed relatively worse (FDiv), or better (FD), than others in both datasets (**Figure 5, Figure 6, Figure A 1, Figure A 2**). Detailed graphs of results for all parameter combinations on both datasets are included in Appendix 1.

**Figure 6.**
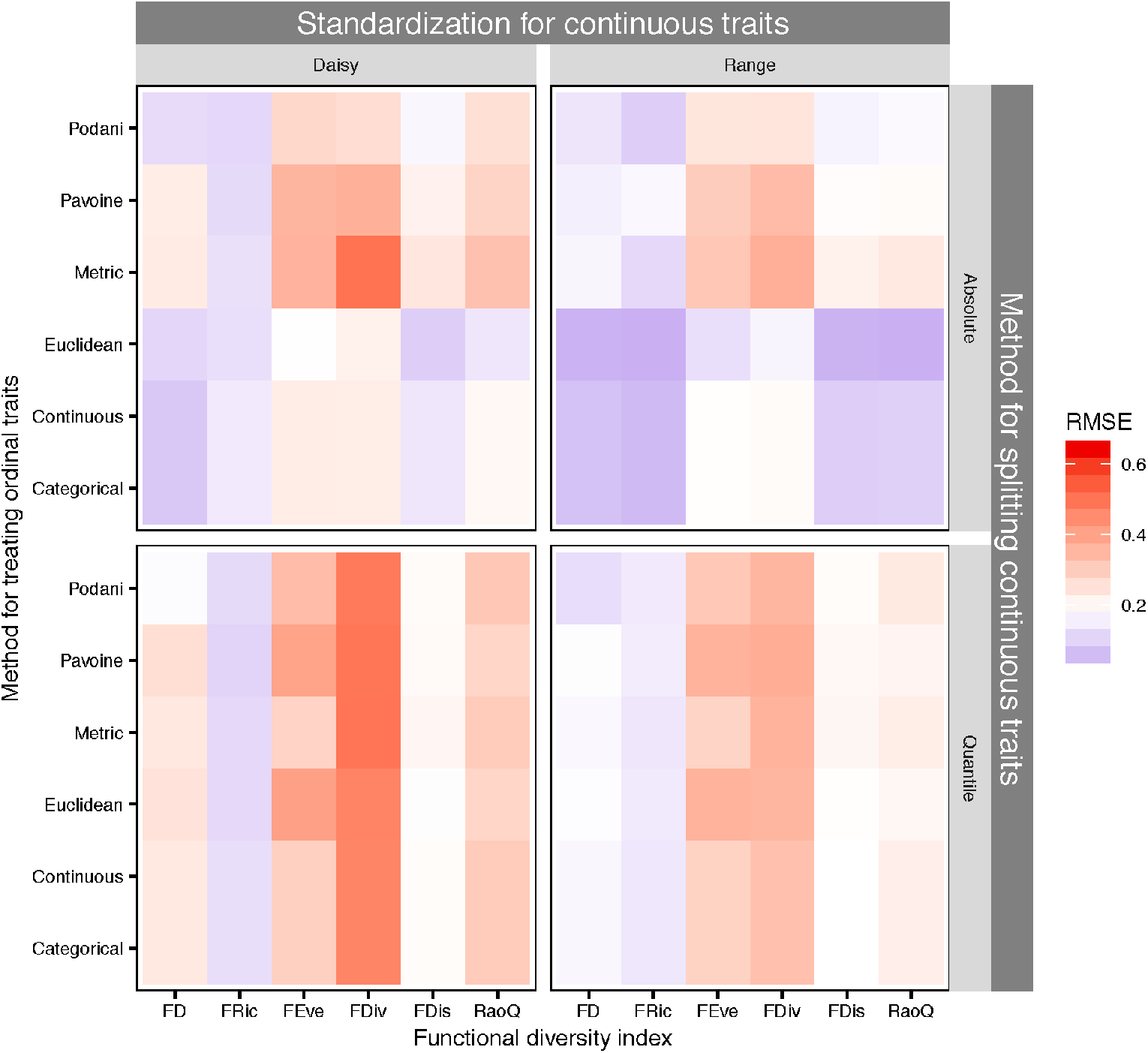
The importance of parameter combinations in determining how closely treatment functional diversity values match original index values in the Tussock dataset. Colors designate the average root mean squared error (RMSE) between the original continuous functional diversity index values and the treatment values obtained from artificially discretized data. Red hues show parameter combinations that give a worse RMSE than the average for all parameter combinations (0.17) and blue hues show parameter combinations that return RMSE better than average.

For ordinal traits using the Podani method of Gower’s distance, a common default parameter combination, discretizing continuous data into more levels increases similarity between original and treatment data as expected, but not monotonically (**Figure 4**). Depending on the underlying distribution of the original continuous data, nine or ten levels may make discretized treatment index values more different than only discretizing the data into three levels. Increasing the number of discretized traits increases error in the RaoQ index value as expected, but surprisingly only if the data have been discretized into seven to ten levels. Data discretized into three to six levels actually produce index values more similar to original continuous values as the number of the traits discretized increased. In the Roadside dataset, different Gower’s distance methods show large differences in RMSE, but only when the original continuous data are discretized using absolute splits (**Figure A 2**). Different Gower’s distance methods produce almost identical RMSE if traits are split along quantiles. And counterintuitively, far from losing information, treating ordinal traits as unranked categories actually approximates the original index values better than almost all other methods (**Figure 5, Figure A 1**).

## Discussion

When comparing values from the treatment data to the original data (**Figure 6, Figure A 2**), certain results are expected. Treatment index values should match original index values standardized by range better than those standardized by the daisy function because the treatment data are already standardized by range with Gower’s distance. We would also expect that increasing the number of traits discretized, or reducing the number of discrete levels, should make treatment index values, on average, more different than original index values. Euclidean distances are likely to return similar values on continuous and discretized traits compared to distances using a more complicated rank interchange formula. Beyond these simple predictions, however, it is hard to see any consistent pattern in the results except that small parameter changes can have large and unpredictable results.

It should not be surprising that using different parameters and different methods for calculating distances between species will produce different functional diversity index values. Different methods represent different mathematical equations and ways of parsing data. Results should be different. What is surprising is how large and unpredictable these differences are, and how much they would impact empirical functional diversity analyses. For example, a researcher following the typical workflow described above with continuous trait data standardized by the daisy function would find that in the Tussock dataset, FDiv decreases with increased fertilizer application up to 250 kg ha-1 yr-1 before increasing again, adjusted R^2^ = 0.59, p < 0.0001 (**Figure 7**, red line). This researcher could then go on to hypothesize why increasing exogenous nutrient loads first drives species closer together in functional space before driving them back apart. The results would surely be of great practical use for preserving ecosystem functions in or near crop-dominated landscapes. However, consider a second researcher. This researcher uses the same data except that two of the eight functional traits, leaf dry matter content and leaf phosphorous concentration, are measured only as “Low”, “Medium”, or “High”. However, using typical workflows and choosing Podani’s (1999) metric method for Gower’s distance on ordinal traits, this researcher will find the exact opposite pattern (**Figure 7**, blue line): FDiv increases up to 250 kg per ha per year and then decreases, adjusted R^2^ = 0.55, p < 0.0001. A third researcher using identical methods and data as Researcher #2, except that leaf nitrogen concentration instead of leaf dry matter content is now “Low”, “Medium”, and “High” will find yet another pattern: FDiv increases linearly with fertilizer loads, adjusted R^2^ = 0.34, p < 0.001 (**Figure 7**, purple line).

**Figure 7.**
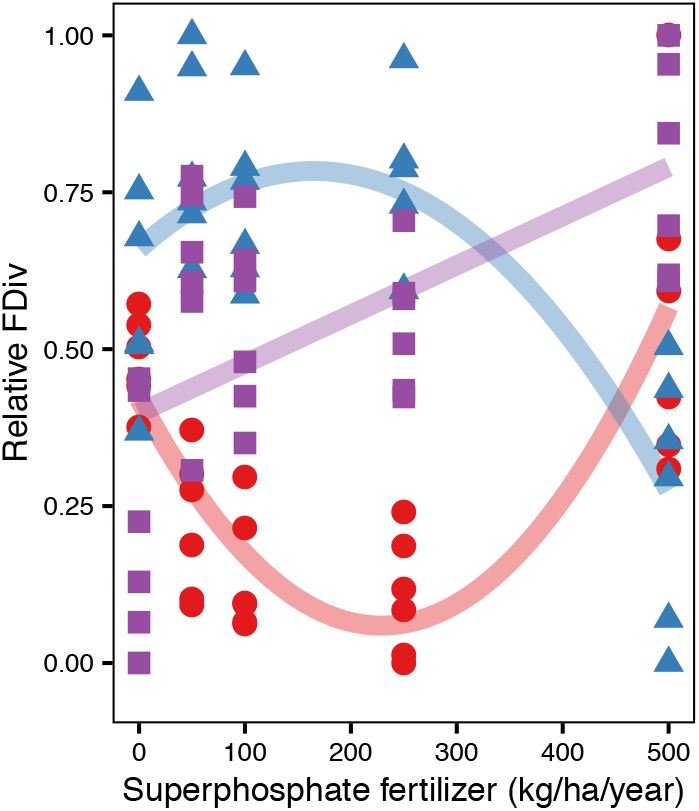
Small changes in methodology can have a large effect on perceived functional diversity gradients. The graph shows the effect of different superphosphate loads on the relative functional divergence (FDiv) of 30 real grassland plots in New Zealand from the Tussock dataset. The red line and points show the relationship if original continuous data, daisy standardization, and Euclidean distances are used. The blue line and points show the relationship if two traits, leaf dry matter content and leaf phosphorous concentration, are treated as ordinal variables, “Low”, “Medium”, and “High” and Gower’s distance (Podani method) is used. The purple line and points show the relationship if all the parameters of the blue line are followed except that leaf nitrogen concentration instead of leaf dry matter content is treated as an ordinal trait, “Low”, “Medium”, and “High”.

FDiv was admittedly the worst performing index considered, but this example using real data and standard methods still raises doubts about the reliability of any functional diversity analysis. If we can fundamentally change a perceived underlying functional diversity relationship by discretizing one trait instead of another, then we have little hope of controlling for the varied effects of different taxa, traits, methods, and study systems to arrive at general principles about functional diversity. Gower’s distance could be avoided by making Euclidean distances on continuous data the gold standard for analysis. But if only continuous data are used, what effect will removing likely non-random subsets of genuinely discrete traits (e.g., C3, C4, and CAM pathways) have on functional diversity analyses? We desperately need more basic research into functional diversity methods and indices before we can begin to make meaningful interpretations of natural patterns.

Of all the treatment parameter combinations, discretizing data at absolute splits and taking Euclidean distance on the resulting ordered levels produced the greatest similarity to the original index values. Although not optimal, using traits with four or more roughly equally spaced ordered levels and Euclidean distance may be a workable stopgap for including discrete characters in functional diversity analyses until we can devise a more robust methodology. Treating ordinal values with Euclidean distance is technically improper mathematically, but all continuous traits are actually discrete at some scale. Consider a researcher measuring plant heights in a population of forbs ranging from around 3 to 95 mm tall; a ruler marked only by centimeters would be a coarse but methodologically sound tool. Using ten ordered levels is operationally no different. For functional diversity indices, Petchey and Gaston’s (2007) modified dendrogram-based FD may be the best choice at present, as it was the least sensitive to parameter changes considered here, and is easily compared to phylogenetic diversity. However, it unrealistically assumes hierarchical functional variation among taxa (Petchey and Gaston 2006) and is known to reflect methodological choices more when communities have similar species richness (Poos et al. 2009).

No methodology will be perfect for all situations, and any methodology must be based on sound statistical and biological arguments (Pavoine et al. 2009) that best suit the ecological question at hand (Lepš et al. 2006). The measurement of distances between species considered here is just one critical step in functional diversity analysis (Pavoine et al. 2009). Others (Lepš et al. 2006, Lavorel et al. 2008) have shown how small methodological choices in data collection can also have outsized impacts on functional diversity index values. New methods will have to acknowledge this uncertainty and be able to integrate across multiple parameter settings to provide more robust measurements of functional diversity.

## Acknowledgements

Thank you to K. Thompson and E. Laliberté for generously providing datasets for simulations and to A. Chandra and K. Nelson for computational support.

---

**Figure A 1.**
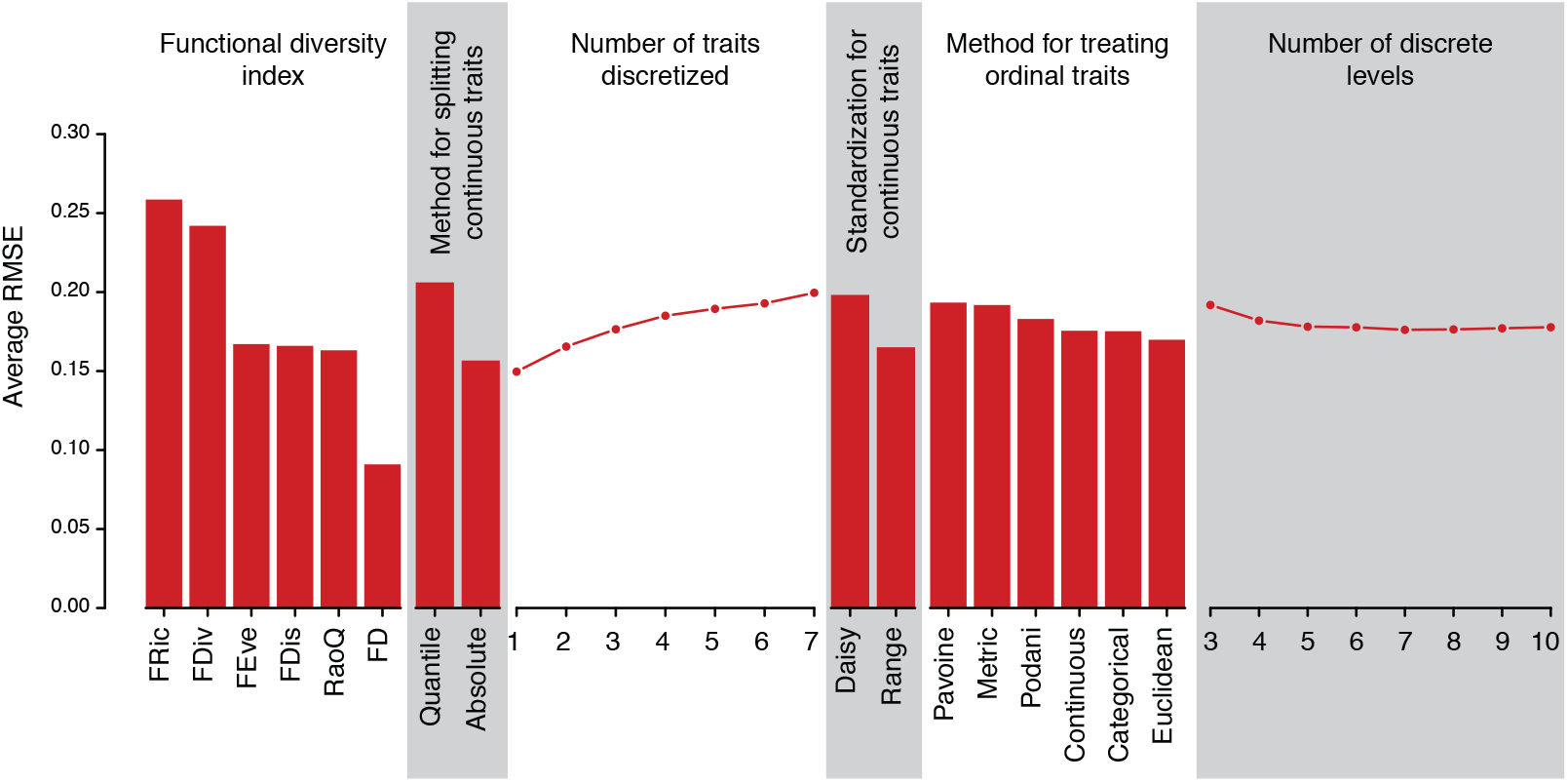
The importance of individual parameters in determining how closely treatment functional diversity values match original index values in the Roadside dataset. The y-axis shows the average root mean squared error (RMSE) between original and treatment values expected from that parameter holding all other parameters constant.

**Figure A 2.**
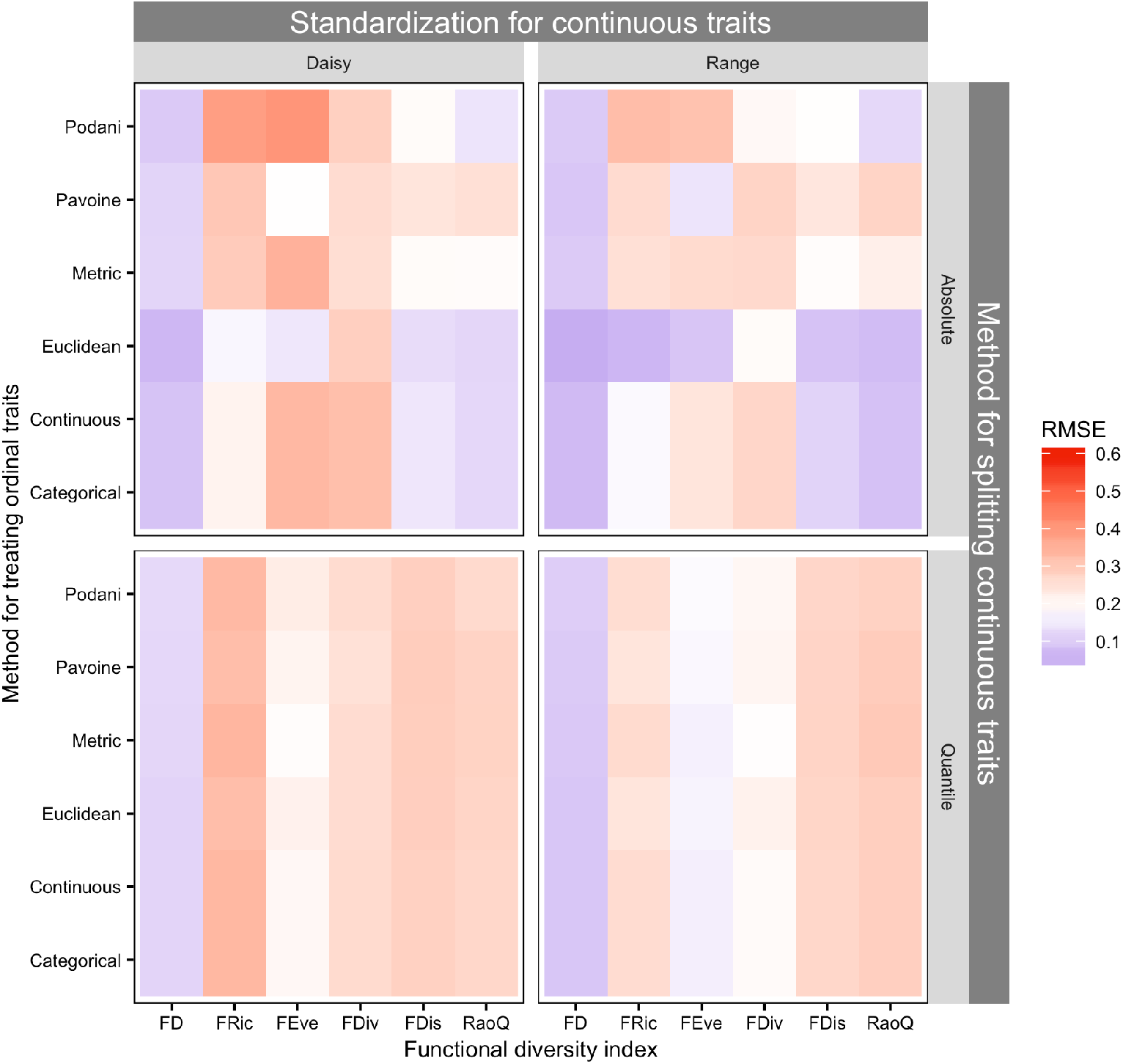
The importance of parameter combinations in determining how closely treatment functional diversity values match original index values in the Roadside dataset. Colors designate the average root mean squared error (RMSE) between the original continuous functional diversity index values and the treatment values obtained from artificially discretized data. Red hues show parameter combinations that give a worse RMSE than the average for all parameter combinations (0.18) and blue hues show parameter combinations that return RMSE better than average.

